# Effects of chronic thermal stress on the reproductive performance of male *Octopus maya*

**DOI:** 10.1101/267476

**Authors:** Laura López-Galindo, Clara Galindo-Sánchez, Alberto Olivares, Omar Hernando Avila-Poveda, Fernando Díaz, Oscar E. Juárez, Fabiola Lafarga, Jordi Pantoja-Pérez, Claudia Caamal-Monsreal, Carlos Rosas

**Affiliations:** Departamento de Biotecnología Marina, Centro de Investigación Científica y Educación Superior de Ensenada, Ensenada, Baja California, México; Facultad de Ciencias del Mar y Recursos Biológicos, Universidad de Antofagasta, Departamento de Biotecnología, Antofagasta, Chile; Facultad de Ciencias del Mar, Universidad Autónoma de Sinaloa, Mazatlán, Sinaloa, México; Dirección de Cátedras-CONACYT, Consejo Nacional de Ciencia y Tecnología (CONACYT), Ciudad de México, México⁴; Departamento de Acuicultura, Centro de Investigación Científica y Educación Superior de Ensenada, Ensenada, Baja California, México; Unidad Multidisciplinaria de Docencia e Investigación, Facultad de Ciencias, Universidad Nacional Autónoma de México, Sisal, Yucatán, México; Laboratorio Nacional de Resiliencia Costera, CONACYT, Sisal, Yucatán, México

**Author notes:** corresponding author (CR). **Summary statement** Temperature affects the physiology and the reproductive performance of male *Octopus maya*, an important fishing resource in the Yucatan Peninsula.

**Keywords:** sperm quality, testis damage, physiological condition, multiple paternity, paternal contribution

## Abstract

In female *Octopus maya* the reproductive success has well-defined thermal limits; beyond which, spawning, number of eggs, fecundity, and the viability of the embryos are reduced. Observations of wild male *O. maya* suggest that temperatures below 27°C favour their reproductive performance. From these observations we hypothesize that, as in females, the temperature modulates the reproductive performance of adult *O. maya* males. The study was directed to evaluate the physiological condition, reproductive success, and histological damage in testis of male *O. maya* exposed to thermal stress, to determine the implications of ocean warming over their reproductive performance. High temperatures (28-30°C) negatively affect the growth and health of male *O. maya.* In octopuses maintained at 30°C, as a consequence of the thermal stress we observed an increment in the haemocytes number, a reduction in the oxygen consumption rate, and an inflammatory process in the testis. The number of spermatozoa per spermatophore was not affected by temperature, but higher spermatophores production was observed at 30°C. The paternity analysis showed that the offspring had multiple paternity with an average of 10 males contributing in a single spawn. The paternal contribution was affected by temperature with high, medium, or no paternal contribution in animals maintained at 24°C (control group), 28°C, and 30°C, respectively. The temperatures from 28°C to 30°C deeply affected the reproductive performance of *Octopus maya* males.

## Introduction

Aquatic environments are thermally heterogeneous in time and space. Organisms inhabiting these environments, specifically ectotherm organisms, show morphological, behavioural, and physiological mechanisms (phenotypic plasticity) that give them adaptive capabilities to cope with environmental changes (Pigliucci, 1996; Somero, 2010; Bozinovic and Pörtner, 2015; Deutchs et al., 2015; Piasečná et al., 2015). Animal physiology, ecology, and evolution are affected by temperature, It is also expected that community structure will be strongly influenced by global warming (Nguyen et al., 2011). For example, temperature seemed to play the most important role in structuring the distribution of cephalopod body size along the continental shelves of the Atlantic Ocean, while resource availability, seasonality, or competition only played a limited role in determining latitudinal body size patterns (Rosa et al., 2012).

In the eastern region of the continental shelf of Yucatan Peninsula (YP), Mexico, a summer upwelling allows sub-superficial subtropical water from the Caribbean (between 150 and 200 m deep) to enter the shelf with temperatures between 16°C and 22°C (Enriquez et al., 2013a). This cold water mass, besides functioning as an external temperature control for the shelf, transports nutrients which are used by primary producers (Enriquez et al., 2010). This upwelling affects only the eastern portion of the YP continental shelf provoking a summer thermal gradient that runs from the western to the eastern shelf from high to low temperatures, offering different environments to aquatic species of the zone (Zavala-Hidalgo et al., 2003; Zavala-Hidalgo et al., 2006; Ciencias de la atmósfera, http://uniatmos.atmosfera.unam.mx/ACDM/).

*Octopus maya* is endemic to the YP continental shelf. This species is the most important octopus fishery in the American continent, with an annual production fluctuating between 8,000 and 20,000 Tons (SAGARPA, 2013; Galindo-Cortés et al., 2014; Gamboa-Álvarez et al., 2015). *O. maya* is an ectotherm organism particularly temperature-sensitive (Noyola et al., 2013a; Noyola et al., 2013b) that can be affected in its morphology, behaviour, physiology and reproduction by changes in ambient temperature with spatio-temporal fluctuations. Predictions of the thermal processes on the YP shelf indicate that sea temperatures may rise between 2.5 to 3°C in the zone where upwelling has no effect (Enriquez et al., 2013b; Saldfvar-Lucio et al., 2015). This temperature increase could be deleterious for this especies, affecting the regional fishing economy. Gamboa-Álvarez et al., (2015) observed that during the August-December fishing season, the greatest abundances of *O. maya* was found along the Campeche coast (western zone, without upwelling influence), where small octopus were fished; whereas, in the eastern zone, less abundances were recorded, but octopus with higher biomass were caught. It was also observed that the *O. maya* wild population reproduces year round in the YP eastern zone, due to low temperatures maintained by the summer upwelling; whereas in the western zone, reproduction occurs only during the winter storms (‘nortes’ season, November-February), when low temperatures favour egg-laying (Avila-Poveda et al., 2015; Markaida et al., 2016; Angeles-Gonzalez et al., 2017).

In laboratory conditions, at 31°C the spawning of female *O. maya* was significantly reduced and only 13% of the total females (n= 32) spawned, while the few fertilized eggs (embryos) were not developed or died after two weeks (Juárez et al., 2015). It was observed that females exposed to a temperature decrease of 1°C every 5 days and starting at 31°C, only 87% spawned after temperatures reached less than 27°C, and of these only 50% of the eggs laid (mean 530 eggs per spawn) were fertilized (Juárez et al., 2015). Those results suggested that temperature could be deleterious to sperm stored in the spermathecae of the oviductal glands, which play a crucial role in octopus reproduction (Olivares et al., 2017). At a later date, the performance of juveniles hatched from those thermal stressed females was evaluated (Juarez et al., 2016). Results obtained in that study showed that juveniles from stressed females had lower growth rate and twice the metabolic rate than hatchlings coming from unstressed females, providing evidence that temperature stress experienced by females has consequences on the performance of hatchlings, with effects on the biomass production and survival.

To date, a small number of studies have investigated multiple paternity within cephalopods by using microsatellite markers demonstrating that multiple paternity could be a common characteristic in octopus species. Voight and Feldheim (2009) sampled *Granelodone boreopacifica* juveniles and found at least two genetically distinct sires that contribute to the progeny. Quinteiro et al., (2011) found evidence of between two to four siring males in egg clutches of *O. vulgaris.* In *Euprymna tasmanica* samples of egg clutches revealed evidence of multiple paternity with two to four sires involved in the contribution to the progeny (Squires et al., 2014). Larson et al., (2015) sampled *Enteroctopus dofleini* eggs finding up to four males contributing to the progeny.

There is enough evidences demonstrating that temperatures higher than 27°C have serious consequences on the reproductive performance and success of female *O. maya.* In this sense, new questions arise: As was observed in females. Is 27°C a thermal threshold for reproductive performance of *O. maya* males? Do *O. maya* males have the physiological mechanisms that allow them to compensate possible damages at temperatures higher than 27°C? To address these questions, we designed a series of experiments to evaluate the effects of fixed temperatures (24°C, 28°C, and 30°C) on adult males of *O. maya* through assessment of their: i) Physiological condition, evaluating the specific growth rate, weight gain, digestive gland index, blood haemocytes and hemocyanin concentration, osmotic capacity, and oxygen consumption; ii) Reproductive performance, evaluated through sperm quality and its relationship with histological characteristics of the testis, and iii) Reproductive success, estimated through the proportion of hatchlings generated by each male in each spawning. Wild adult females were mated with laboratory stressed males. Considering that multiple paternity can be present in *O. maya,* a paternity analysis implementing specific microsatellite markers was performed to assess the reproductive success of the experimental males.

To our knowledge, this is the first work that investigates the chronic thermal effect in the reproductive performance and success of male octopuses.

## Material & methods

### Ethics Statement

In this study, octopuses were anesthetized with ethanol 3% in seawater at experimental temperatures (Estefanell et al., 2011; Gleadall, 2013) to induce narcotisation to enable humane killing (Andrews et al., 2013) in consideration of ethical protocols (Mather and Anderson, 2007), and the animal’s welfare during manipulations (Moltschaniwskyj et al., 2007). Our protocols were approved by the experimental Animal Ethics Committee of the Faculty of Chemistry at Universidad Nacional Autonoma de México (Permit number: 0ficio/FQ/CICUAL/099/15). We encouraged the effort to minimize animals stress and the killing of the minimum necessary number of animals for this study.

### Animal Capture and laboratory conditioning

Seventy-two *O. maya* adult males with body weight above 300 g were captured in the Sisal coast of the Yucatan Peninsula (21°9′55″N, 90°1′50″W), by using the local drift-fishing method known as “Gareteo” (Solís-Ramírez, 1967; Pascual et al., 2011). Male octopuses were caught during three collection trips from June to September of 2015. All captured males above 300 g were anatomically mature with a developed reproductive system, thus sexually mature (Avila-Poveda et al., 2016). Octopuses were maintained in a 400-L black circular tank with seawater recirculation and exchange during the capture and then transported to the Experimental Cephalopod Production Unit at the Multidisciplinary Unit for Teaching and Research (UMDI-UNAM), Sisal, Yucatan, Mexico. Octopuses were acclimated for 10 d in 6 m diameter outdoor ponds provided with aerated natural seawater (26 ± 1°C). The ponds were covered with black mesh reducing direct sunlight to 70%, and connected to seawater recirculation systems coupled to protein skimmers and 50 μmb bag filters. PVC 50 mm diameter open tubes were offered as refuges in proportion 2:1 per animal. Octopuses were fed individually twice a day with a paste made with squid and crab meat at ratio of 8% of its body weight (Tercero et al., 2015).

### Experimental design

After the conditioning period the 72 adult male *O. maya* were randomly distributed in 80 L individual tanks at three different temperatures, 24, 28, and 30°C with n=23 specimens per treatment, and mean weights of 584 ± 193 g ww, 692 ± 203 g ww, and 557 ± 160 g ww, respectively; P < 0.05. Males were maintained in experimental conditions during 30 d and feed with the same paste used during the conditioning period. Seawater in tanks was maintained in a semi-closed recirculation system coupled with a rapid-rate sand filter and 36 ± 1 ppt salinity, dissolved oxygen higher than 5 mg L^−1^, pH above 8, photoperiod of 12L/12D and a light intensity of 30 Lux cm^−2^. For the experimental temperatures above 26°C, seawater temperature was gradually increasing 2°C per day until the experimental temperature was reached. Temperatures of 28°C and 30°C were controlled with 1,800-Watt heaters connected to automatic temperature controllers, while temperature of 24°C was controlled with a titanium chiller and the air conditioning of the experimental room.

### Physiological condition

#### Specific growth rate and digestive gland index

We used 23 octopus adult males to evaluate physiological condition of animals exposed to experimental treatments. These animals were classified as PREmating, taking into account that they were only exposed to experimental temperatures for 30 d. Before measurements, animals were anesthetized with alcohol 3% in sea water at the actual experimental temperature; this procedure took 3-6 min. The organisms were considered anesthetized when the respiration was imperceptible (Gleadall, 2013). Afterwars, each octopus was weight and a blood sample of 100 to 150 μL was drawn using a catheter inserted in the dorsal aorta. The sample was kept in ice until the haemocytes count. Once samples were obtained, octopus were euthanized cutting the brain in the middle of the eyes (Gleadall, 2013). Afterwars, the reproductive system and total digestive gland were extracted.

Total weight gain (WG) is the difference between the octopuses’ wet weight at the beginning and the end of the experiment. Specific growth rate (SGR) was calculated as SGR = [(LnWf - LnWi) / t] ∗ 100, where Wf and Wi are the octopuses’ final and initial wet weights, respectively, Ln is the natural logarithm and t is the number of experimental days. Survival was calculated as the difference between the number of animals at the beginning and at the end of the experiment. The Digestive gland index was calculated as: DGI= (DGW / Wf)*100: where DGW= digestive gland weight in g (Cerezo-Valverde et al., 2008).

#### Total haemocytes count and hemocyanin concentration (Hc)

Total haemocytes count (THC) was determined by processing the 10 μl of hemolymph sample immediately after extraction. The hemolymph sample was placed in TC10 counting slides with dual chambers and the readings were performed with a TC10™ automated cell counter (Bio-Rad). The hemocyanin concentration was measured by using 990 μl of TRIS 0.1 M (pH 8.0) and 10 μl of hemolymph. These procedures were triplicated. Hemocyanin measurements were performed using a spectrophotometer Genesys 10 with UV lamp (Thermo Scientific) in 1 ml UV cells at 335 nm of absorbance. The Hc concentration was calculated as: Hc = (mean Abs/ɛ)/DF; where Abs = absorbance at 335 nm, ɛ = extinction coefficient (17.26), and DF = dilution factor.

#### Osmoregulatory capacity (OsmC)

The osmotic pressure (OP) of 20 μL hemolymph samples were measured for every octopus in each treatment concurrently with the OP of three water samples in each treatment. OP was measured in a Micro osmometer 3MoPLUS (Advanced Instruments). The osmotic capacity was calculated as: OsmC= hOp-wOp; where hOp= hemolymph osmotic pressure and wOp= water osmotic pressure.

#### Oxygen consumption (VO2)

The oxygen consumption (VO2) was measured using a continuous flow respirometer where respirometric chambers were connected to a well-aerated, recirculating seawater system (Rosas et al., 2008). Eight male octopi per experimental condition were placed in 15 L chambers with an approximate flow rate of 5 L min^−1^. All animals were allowed to acclimate to the chambers for 30 min before measurements were made. A chamber without an octopus was used as a control. Measurements of dissolved oxygen (DO) were recorded for each chamber (at entrance and exit) every minute during 4 h using oxygen sensors attached to flow cells, which were connected by an optical fibre to an Oxy 10 mini-amplifier (PreSens©, Germany). The sensors were calibrated for each experimental temperature using saturated seawater (100% DO) and a 5% sodium sulphate solution (0% DO).

The oxygen consumption (VO_2_) was calculated as VO_2_= [(O_2i_-O_2o_) *F] / Bw; where O_2i_= oxygen concentration of the water inlet (mg/L^−1^), O_2o_= oxygen concentration of the water outlet in each experimental chamber (mg/L^−1^), F= water flow rate (L/h^−1^), BW= octopus total body weight (g).

### Reproductive performance

#### Reproductive indexes and sperm quality

To establish the sexual maturity and reproductive activity of the experimental octopuses during 30 d of thermal exposure, the following indexes were estimated:

The Gonadosomatic index, GSI= (TW/BW)*100; Spermatophoric complex index: SCI= (SCW/BW)*100; Maturity coefficient: MCO= [(TW+SCW)/BW]*100; where TW= testis weight (g); SCW= spermatophoric complex weight (g); BW= total body weight (g) (Krstulovic-Sifner and Vrgoc, 2009; Sivashanthini et al., 2010; Rodrigues et al., 2011).

The total number of spermatophores (STN) for each Needham’s sac was counted. Three spermatophores per octopus were taken to evaluate the total number of spermatozoa (TSC), as well as the number and percentage of alive (TASC and ASP) and dead spermatozoa for each experimental treatment. Spermatophores were homogenized in 2 ml of Ca^2^+ free solution. Then 10 μl of the homogenate was mixed with 4% tripan blue (v/v). Readings were performed in a TC10 Automated Cell Counter (Bio-Rad) with 10 μl of the mix.

#### Testis Histology

A portion of the gonad of approximately 1 cm^3^ was taken by performing a perpendicular cut to the tunica albuginea (the fibrous connective membrane that covers the testis, “testis wall”). That portion of the gonad was fixed in Davidson’s fixative for 3 d (Elston, 1990), rinsed in 70% ethanol, dehydrated in an ethanol series, cleared in Ultraclear^®^, permeated and embedded in Paraplast^®^ tissue embedding medium (m.p. 56°C). Sections of 5 μm were stained with Harris’s Hematoxylin and Eosin regressive method (Howard et al., 2004). Slide examinations were performed at 400x and digital images were obtained with a digital imaging system (Micrometrics^®^ SE Premium 4.4 software, ACCU_SCOPE) mounted on an Olympus H30 compound microscope.

Twenty seminiferous tubules (ST) close to the tunica albuginea found in longitudinal sections were randomly selected and two widths and three heights were measured: width of ST and lumen, height of the strata of germ cells (SGC: spermatogonia, spermatocytes of 1^st^ and 2^nd^ order, round spermatid, ovoid spermatid, and elongated spermatid), height of proliferative stratum (PS: spermatogonia, spermatocytes of 1^st^ and 2^nd^ order, and round spermatid), and height of differentiation stratum (DS: ovoid spermatid and elongated spermatid), where SGC = PS + DS (Fig. S1). The total relative surface area measured was then considered to the nearest 5 mm^2^. The percentage of disorders in the area of germinal cells such as completely acidophilic bodies, or with basophilic material, and vacuolated basal compartments were calculated.

### Male Reproductive success

#### Mating Protocol

Six of the 23 octopuses for each experimental temperature were mated with two females 2:1 that were previously maintained at 24°C for 20-30 d. The *Octopus maya* females were maintained in 80L natural seawater tanks in similar conditions that males, but at a 24°C constant temperature. This experiment was done trying to ensure that each female was mated with at least three different males from the same experimental temperature (Fig. S2). Male octopuses were placed in the female tanks and acclimated during 30 min. Males were allowed to mate during 4 to 6 h and then returned to their experimental tank. Males used in the mating protocol were sacrificed 12 h after mating following the protocol previously described. Those males were considered POST-mated and classified as POST.

Pregnant females were maintained in individual tanks until spawning, and fed twice a day. After spawning, wet weight was recorded. Each spawning was placed in an artificial incubator (Rosas et al., 2014) during 45–50 d, with a range temperature of 22°C to 24°C, and constant salinity, pH, aeration, and seawater recirculation. Data of the number of eggs per spawn, number of hatchlings, hatchlings wet weight, deformities, fecundity, and survival of hatchlings after 10 d fasting were recorded. To evaluate the quality of hatchlings obtained from females mated with males exposed at different experimental temperatures, hatchlings survival was evaluated by placing 20 juveniles in PVC tubes individualized without feeding during 10 d (Rosas et al., 2014).

### Statistical analyses

Data were expressed as mean ± SD. Differences among values of each measurement (widths and heights) throughout the treatments (temperature and condition PRE-POST) were evaluated by two-way ANOVA followed by Fisher LSD (least significant difference) tests. Data transformation were applied to obtain normality and homocedasticity to fulfill the ANOVA assumptions (McCune et al., 2002; Zar, 2010). Statistical analyses were carried out using STATISTICA7^®^ (StatSoft). Statistical significance was accepted if *P* < 0.05.

No significant differences were found between the PRE and POST reproductive conditions among all tested parameters; therefore, the data of the 23 tested octopuses were used to calculate the mean for the different parameters and only thermal exposure was considered as the main effect factor.

### Paternity analyses

#### DNA extraction

The DNA of 47 hatchlings per spawn, for a total of 282, and breeders, six females and 17 males, was extracted from arm tissues. Approximately 30 mg of tissue were homogenized with mortar and pestle, adding liquid nitrogen. DNA was extracted using the DNeasy^®^ Blood and Tissue kit (Qiagen) following the supplier instructions. The concentration and purity of each DNA sample were measured with a Nanodrop (Thermo-Scientific) spectrophotometer. The DNA integrity was assessed with an electrophoresis in agarose gel (1%) at 85V for 40 min.

#### Microsatellite amplification

To obtain the hatchlings and breeders genotype, five polymorphic microsatellite loci previously characterized (Juárez et al., 2013; Table 1) were selected for polymerase chain reaction (PCR) amplification. PCR primers were marked with 6FAM, VIC, PET, and NED fluorescent dyes (Applied Biosystems) for subsequent fragment analysis. The PCR for each microsatellite was performed in a thermal cycler CFX96 Touch™ (Bio-Rad), on 96-well plastic wells. The 15 μL reaction volumes contained: 3 μL Buffer (5X), 0.9-1.5 μL MgCl_2_ (25 mM), 0.3 μL dNTP (10 mM), 0.15 μL of each primer (10 μM), 1 μL DNA (40 ng/μL), 8.3-9.425 μL H_2_O depending on each locus (specific PCR conditions of each locus in Table 1), and 0.075 μL of Go Taq Flexi DNA polymerase (5 u/μL, Promega).

The general amplification program was: 2 min at 94°C; followed by 35 cycles of 30 sec at 93°C, specific alignment time at specific Tm (Table 1), and 30 sec at 72°C; finally an elongation step was added (10 min at 72°C). Positive and negative controls were included in each plate. The PCR amplicons were verified by electrophoreses in agarose gels (1.5%) at 85V for 40 min. The amplicons marked with different fluorophores obtained from the same sample, were multiplexed for fragment analysis in an AB genetic analyzer (Applied Biosystems).

**Table 1.**
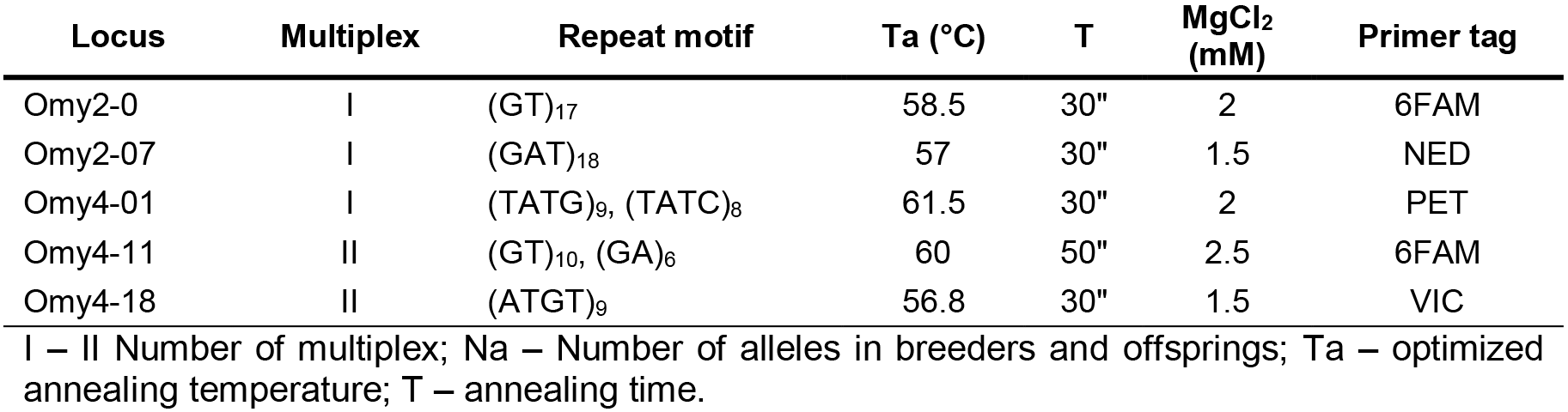
Primer sequences, characteristics and PCR conditions for amplification of 5 microsatellite loci *of O. maya* (Juárez et al., 2013).

#### Fragment Analyses and Genotyping

Fragment analyses were performed in the AB 3730xl genetic analyzer (Applied Biosystems) at the Illinois University Roy J. Carver Biotechnology Center (USA). The allele size in each sample was assigned using the PEAK SCANNER software (Applied Biosystems). The multilocus genotype of each sample (offsprings and breeders) was registered to build a data matrix.

#### Parentage and Data Analyses

The paternity analyses were conducted using two different softwares COLONY 2.0.6.3, and GERUD 2.0. COLONY estimates the maximum number of sires in the spawn using a maximum-likelihood method to assign parentage and sibship groups, if the potential fathers were not sampled the program reconstructs the genotypes (Jones and Wang, 2010). For each spawn, the potential father’s genotypes were inferred, providing the mother, the candidate fathers, and offsprings genotypes as input data for the analysis. If the genotypes of the candidate males did not appear in the inferred father genotypes (paternity), it was assumed that the father was a wild male octopus. GERUD determines the minimum number of paternal genotypes that are necessary to produce the genotypes of the progeny in the spawn based on the Mendelian segregation laws, and the allele frequencies in the spawns, considering consistent maternal genotypes (Jones, 2005). For each spawn, the maternal and offsprings genotypes were used as input for the analysis. Five microsatellite loci were used in the analysis; in some cases loci with missing data were discarded. A correlation between the number of inferred fathers and the experimental conditions was performed.

Observed and expected heterozygosity (H_o_ and H_e_, respectively) of breeders and offsprings, Hardy–Weinberg equilibrium (H_W-E_), and inbreeding coefficient (F_IS_) were obtained using ARLEQUIN 3.5.2.2 software (Excoffier et al., 2005). The FIS index was estimated using the analysis of molecular variance (AMOVA) with 1000 permutations. The number of alleles and allele frequencies (Table S1) were obtained with the ARLEQUIN software.

## Results

### Physiological condition

Total weight gain (WG) and SGR (% d^−1^) were affected by temperature (Table 2; P < 0.05). Total WG of animals maintained at 24 and 28°C were 9 times higher than the observed in octopuses maintained at 30°C. In consequence a SGR 6 times higher was obtained in animals maintained at 24 and 28°C than those maintained at 30°C (Table 2). We observed that octopuses exposed to 30°C not only lost weight but also reduced their food ingest intermittently during the 30 d exposure period. The temperature also affected the DGI (Table 2). The DGI of animals maintained at 24°C was 58% higher than those obtained in octopuses exposed at 28 and 30°C (Table 2; P < 0.05).

Blood parameters were also affected by temperature. A higher concentration of THC was recorded in octopuses exposed to 30°C (2.5×10^6^ ± 1.5×10^6^ cells/ml) in comparison to organisms maintained at 24 and 28°C (Table 2; P < 0.05). The Hc was significantly lower (P < 0.05) at 28°C (1.84 mmol/L) than that observed in animals maintained at 24°C and 30°C (2.10 and 2.27 mmol/L; Table 2; P < 0. 05). Considering that there were no statistical differences between OsmC values obtained in experimental animals, a mean value of 415 + 85 mOsm kg^−1^ was calculated (Table 2; P > 0.05). Temperature affected the routine metabolism of male *O. maya* with values 42% lower in animals maintained at 30°C (0.02 mg O_2_ h^−1^ g^−1^) than those observed in animals maintained at 24 or 28°C (0.03 mg O_2_ h^−1^ g^−1^; Table 2; P < 0.05).

**Table 2.**
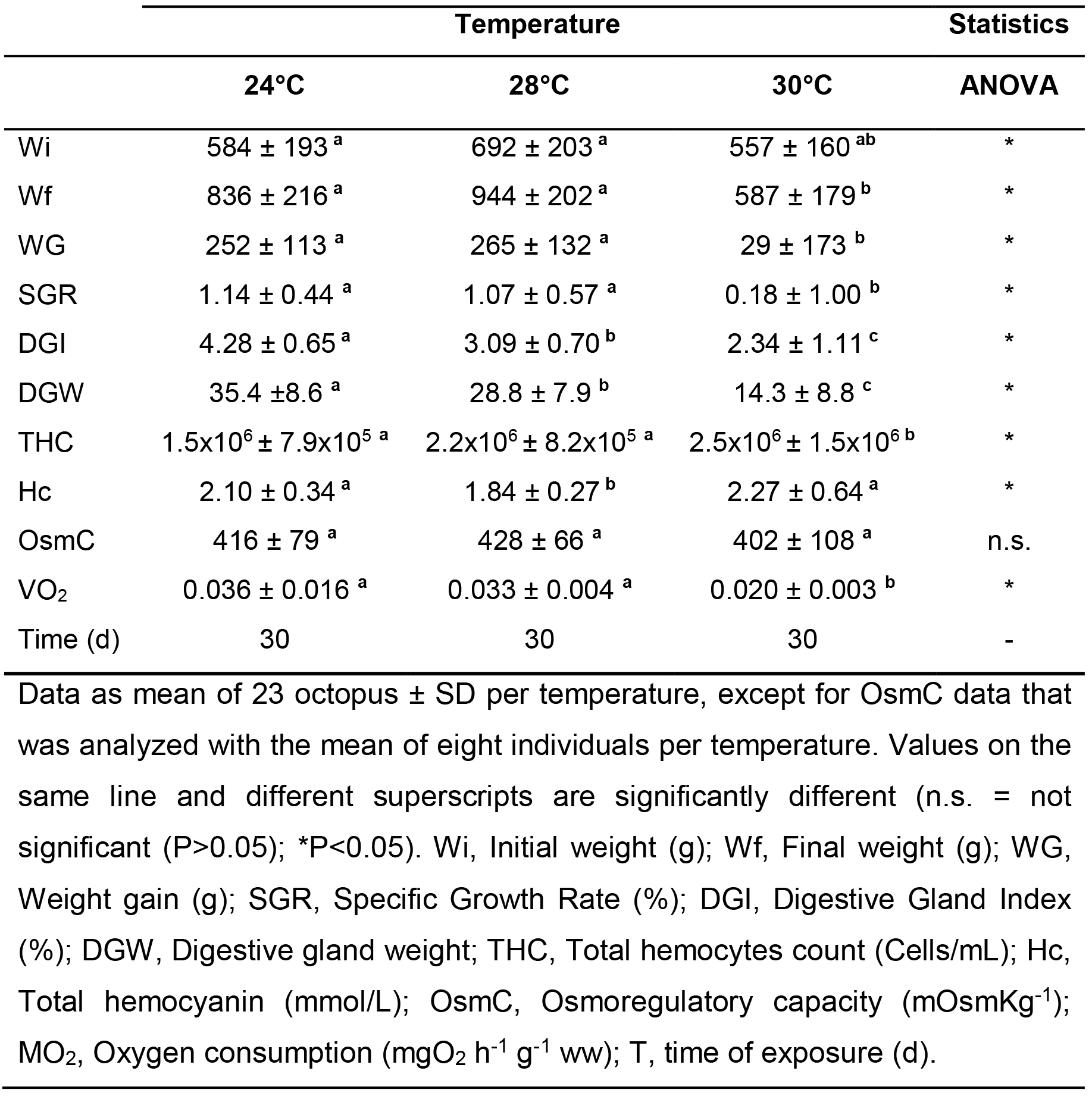
Physiological condition of *O. maya* males exposed to chronic thermal stress.

### Reproductive performance

Temperature did not affect the spermatozoa content per spermatophore (TSC, TASC, and ASP, Table 3; P > 0.05). In contrast an increment of the STN-PRE with temperature was detected with lower values in animals maintained at 24°C (84 spermatophores animal^−1^) than those observed in octopuses exposed to 28°C or 30°C (mean value 129 spermatophores animal^−1^; P < 0.05). The STN-POST also was affected by temperature with low values in animals maintained at 24°C and 28°C (mean value 52 spermatophores animal^−1^) than those observed in octopuses maintained at 30°C (108 spermatophores animal^−1^; Table 3; P < 0.05).

**Table 3.** Reproductive performance and sperm quality indicators calculated for *O. maya* males exposed to chronic thermal stress.

|  | Temperature |  |  | Statistics |
| --- | --- | --- | --- | --- |
|  | 24°C | 28°C | 30°C | ANOVA |
| N | 23 | 23 | 23 | - |
| TSC | $1.3 \times 10^6 \pm 5.7 \times 10^5$ <sup>a</sup> | $1.4 \times 10^6 \pm 4.7 \times 10^5$ <sup>a</sup> | $1.4 \times 10^6 \pm 3.5 \times 10^5$ <sup>a</sup> | n.s. |
| TASC | $5.2 \times 10^5 \pm 2.3 \times 10^5$ <sup>a</sup> | $5.5 \times 10^5 \pm 1.9 \times 10^5$ <sup>a</sup> | $5.3 \times 10^5 \pm 1.3 \times 10^5$ <sup>a</sup> | n.s. |
| ASP | $37.4 \pm 3.4$ <sup>a</sup> | $36.7 \pm 4.9$ <sup>a</sup> | $37.1 \pm 3.8$ <sup>a</sup> | n.s. |
| STN<sup>1</sup> | $84 \pm 20$ <sup>a</sup> | $115 \pm 30$ <sup>b</sup> | $142 \pm 63$ <sup>b</sup> | * |
| STN<sup>2</sup> | $54 \pm 32$ <sup>a</sup> | $49 \pm 17$ <sup>a</sup> | $108 \pm 40$ <sup>b</sup> | * |
| TW | $7.66 \pm 1.93$ <sup>a</sup> | $7.81 \pm 1.97$ <sup>a</sup> | $6.34 \pm 2.72$ <sup>a</sup> | n.s. |
| SCW | $6.70 \pm 3.09$ <sup>a</sup> | $7.63 \pm 1.99$ <sup>a</sup> | $7.87 \pm 2.10$ <sup>a</sup> | n.s. |
| GSI | $0.93 \pm 0.15$ <sup>a</sup> | $0.84 \pm 0.19$ <sup>a</sup> | $1.11 \pm 0.51$ <sup>b</sup> | * |
| SCI | $0.78 \pm 0.22$ <sup>a</sup> | $0.83 \pm 0.24$ <sup>a</sup> | $1.46 \pm 0.60$ <sup>b</sup> | * |
| MC | $1.70 \pm 0.32$ <sup>a</sup> | $1.66 \pm 0.32$ <sup>a</sup> | $2.57 \pm 0.91$ <sup>b</sup> | * |

Values on the same line and different superscripts are significantly different (- not applicable; n.s. = not significant ( $P > 0.05$ ); \* $P < 0.05$ ). N = number of tested octopus; TSC, Total sperm count (cells ml<sup>-1</sup> spermatophore<sup>-1</sup>); TASC, Total alive sperm count (cells ml<sup>-1</sup> spermatophore<sup>-1</sup>); ASP = Alive sperm percentage (%); STN, Spermatophores total number (<sup>1</sup>-PRE, <sup>2</sup>-POST); TW, Testis weight (g); SCW, Spermatophoric complex weight (g); GSI, Gonadosomatic index (%); SCI, Spermatophoric complex index (%); MC, Maturity coefficient (%).

The testis and the spermatophoric complex mean weights (TW and SCW) were not affected by temperature; mean values of 7.3 and 7.4 g animal^−1^ can be calculated for male *O. maya* sampled in this study (P > 0.05; Table 3). The GSI, SCI, and MCO were affected by experimental temperature with significantly higher values in animals maintained at 30°C than observed in octopuses exposed at 24°C and 28°C (Table 3; P < 0.05).

With increasing temperature, a dilation of the seminiferous tubules and their lumen were evident, from 24°C to 28°C increasing 50-60 microns, while from 28°C to 30°C the dilation increased another 80-100 microns. Despite the expansion of the seminiferous tubules and lumen, each one of the two strata (proliferative and differentiation) forming the area of germ cells showed no significant change in height with increasing temperature (P > 0.05), except at 30°C where shrinkage of about 20 microns was observed, mostly the proliferative stratum (spermatogonia, spermatocytes of 1^st^ and 2^nd^ order, and round spermatid). All treatments showed completely acidophilic bodies in all strata of germ cells in an order of 3% to 5%, except octopuses treated at 30°C, which showed a 4-fold of these completely acidophilic bodies compared to the other treatments (Fig. 1). At 30°C we observed acidophilic bodies with basophilic material, and vacuolated basal compartments (Fig. 2).

**Fig. 1.**
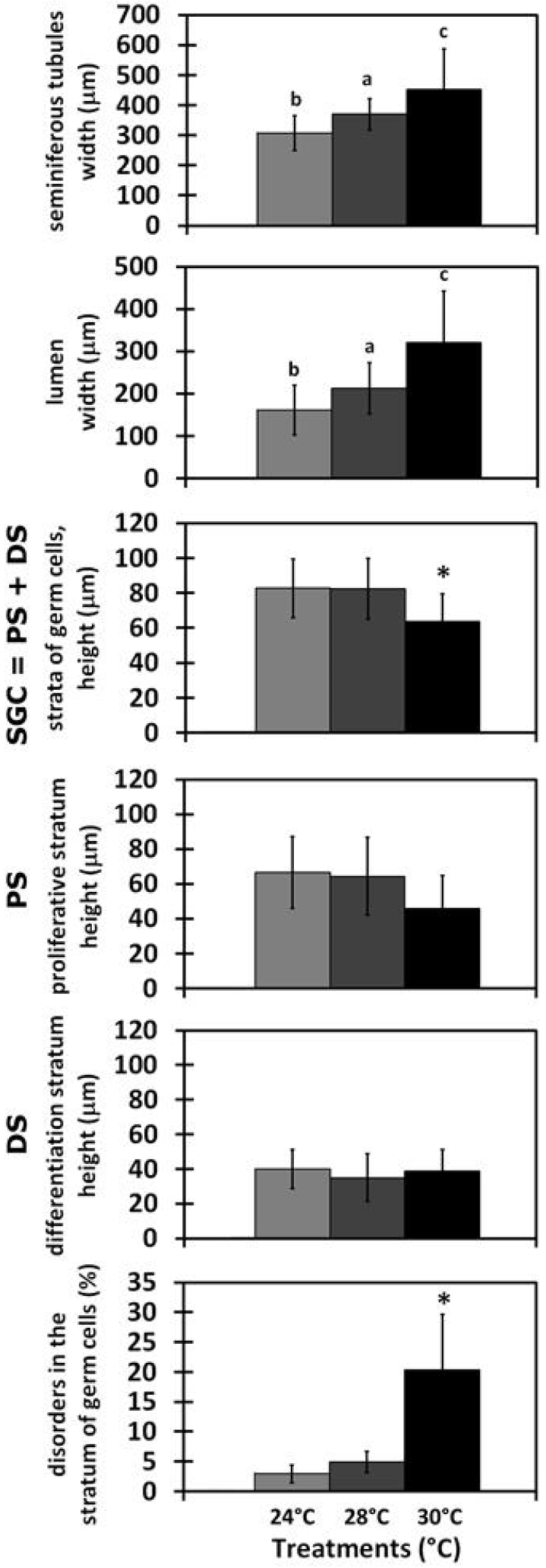
Morphological changes in the germ cells strata (SGC = PS + DS), and the seminiferous tubules lumen during experimental thermal stress. Values are mean ± SD. Different letters indicate significant differences among treatments and asterisks denote significant differences from all other treatments at P < 0.05.

**Fig. 2.**
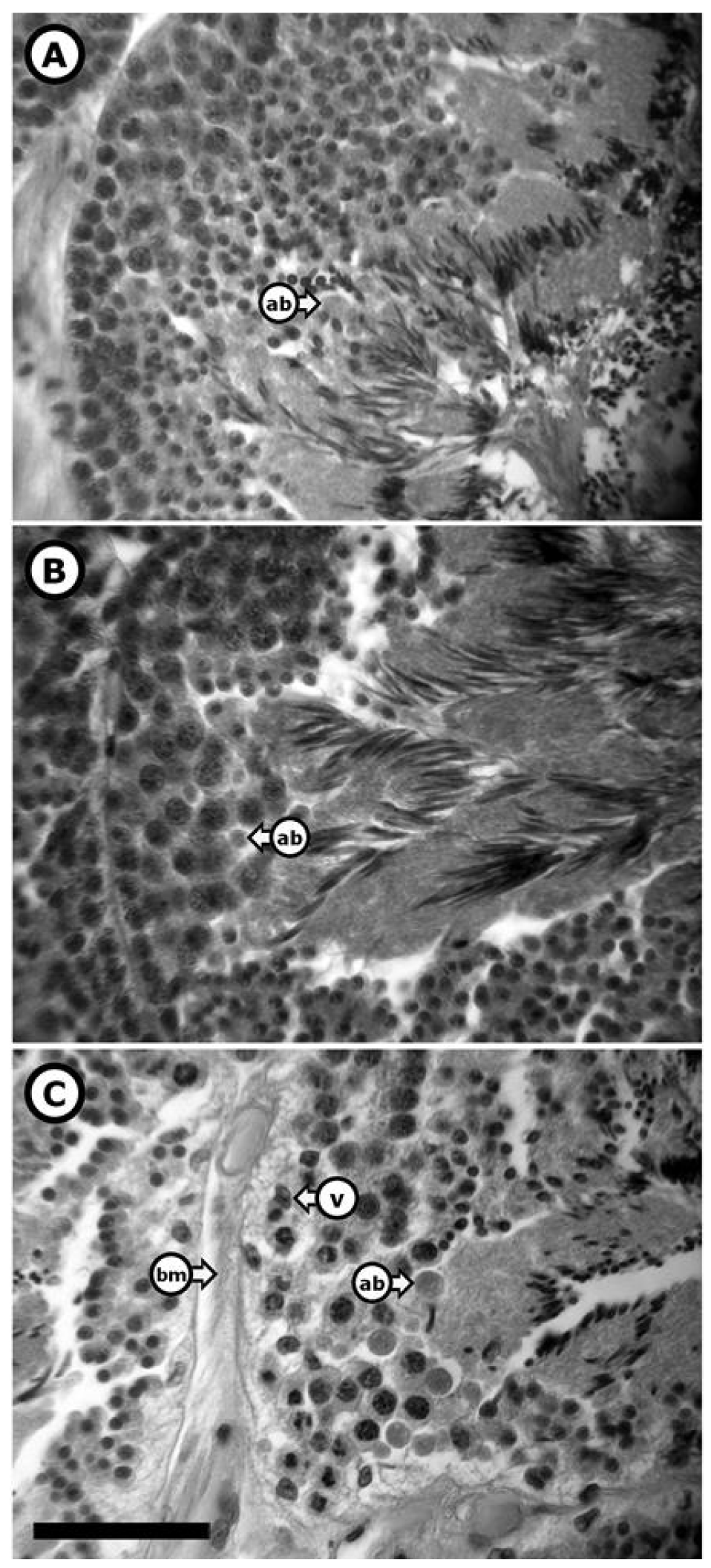
Cross sections photomicrographs of *Octopus maya* seminiferous tubules during the chronic thermal stress. Treatments are: A) 24°C, B) 28°C and C) 30°C. Abbreviations: ab-acidophilic bodies, bm-basement membrane of the seminiferous tubule, v-vacuole in the basal area. General structure followed scheme from figure 2. Scale bars are 50 μm.

### Male reproductive success

#### Fertilization

Fertilization rate was apparently not affected by temperature. All the females mated with males from experimental temperatures spawned normal eggs that developed as embryos and hatched without deformities. Egg fertilization fluctuated between 53% and 92% with no apparent relationship with the experimental temperature experienced by males (Table 4). Also, hatchlings survival after the 10 d fasting was high with percentages that oscillated between 85% and 100%.

**Table 4.** Reproductive capacity of *O. maya* males exposed at different experimental temperatures for 30 d.

|  | Males temperature (°C) |  |  |
| --- | --- | --- | --- |
|  | 24°C | 28°C | 30°C |
| Male Wet weight, g | 699 ± 53 | 780 ± 160 | 519 ± 143 |
| **Reproductive success after mate:** |  |  |  |
| **Female 1** |  |  |  |
| Spawn ID | S24-1 | S28-1 | S30-1 |
| Female Wet weight, g | 597 | 790 | 787 |
| Mating time, h | 4 | 5 | 6 |
| Eggs per spawn | 737 | 797 | 446 |
| Number of hatchlings | 537 | 509 | 411 |
| Hatchlings wet weight, g | 0.11 ± 0.02 | 0.12 ± 0.01 | 0.11 ± 0.01 |
| Survival after 10 d fasting, %b | 90 | 95 | 100 |
| Deformities | 0 | 1 | 2 |
| Fecundity, % | 73 | 64 | 92 |
| **Female 2** |  |  |  |
| Spawn ID | S24-2 | S28-2 | S30-2 |
| Wet weight, g | 628 | 1047 | 553 |
| Mating time, h | 4 | 5 | 6 |
| Eggs per spawn | 772 | 782 | 481 |
| Number of hatchlings | 518 | 418 | 424 |
| Hatchlings wet weight, g | 0.13 ± 0.01 | 0.11 ± 0.01 | 0.09 ± 0.01 |
| Survival after 10 d fasting, %b | 90 | 90 | 85 |
| Deformities | 8 | 1 | 0 |
| Fecundity, % | 67 | 53 | 88 |

Females maintained at temperature 24°C
Mated males: 6 per treatment; Mated females: 2 per treatment; Mated males per female (N=3); Number of hatchlings weighed: 50 per female; Number of hatchlings to evaluate survival after 10 d fasting period: 20 per female.

### Paternity analyses

All the microsatellite loci used in this study were polymorphic and correctly amplified in all samples, showing a high level of genetic diversity (Table 5). Fifty one alleles were detected from 267 individuals (B- Breeder, O- Offspring). N_a_ ranged from four to 9 in B and 6 to 12 in O per locus. H_o_ ranged from 0.35 to 0.83 in B and 0.42 to 0.77 in O, respectively; and H_e_ ranged from 0.43 to 0.85 in B and 0.42 to 0.81 in O, respectively. F_IS_ ranged from -0.187 to 0.202 and 0.009 to 0.075 in B and O, respectively. F_IS_ averages were 0.007 with a P-value of 0. 484 in B and 0.037 with a P-value of 0.014 in O, as a whole. H_W-E_ performed among 10 locus for breeders-offsprings combinations, revealed a significant deviation at four loci (P < 0.05). These four loci were Omy2-0, Omy2-07, Omy4-01, and Omy4-11 in O, while B were within H_W-E_. In the case of Omy4-18 were within H_W-E_ in B and O (Table 5).

**Table 5.** Summary statistics of five microsatellite markers in *O. maya* males.

| Locus | N<sub>a</sub> | H<sub>o</sub> | H<sub>e</sub> | P<sub>H-W</sub> | F<sub>IS</sub> |
| --- | --- | --- | --- | --- | --- |
| Omy2-0 |  |  |  |  |  |
| B | 9 | 0.74 | 0.85 | 0.367 | 0.128 |
| O | 12 | 0.77 | 0.81 | 0.000** | 0.053 |
| Omy2-07 |  |  |  |  |  |
| B | 4 | 0.35 | 0.43 | 0.258 | 0.202 |
| O | 12 | 0.51 | 0.52 | 0.000** | 0.009 |
| Omy4-01 |  |  |  |  |  |
| B | 9 | 0.83 | 0.74 | 0.542 | -0.115 |
| O | 12 | 0.69 | 0.71 | 0.000** | 0.018 |
| Omy4-11 |  |  |  |  |  |
| B | 6 | 0.78 | 0.66 | 0.792 | -0.187 |
| O | 6 | 0.60 | 0.65 | 0.004* | 0.075 |
| Omy4-18 |  |  |  |  |  |
| B | 4 | 0.48 | 0.51 | 0.669 | 0.064 |
| O | 7 | 0.42 | 0.42 | 0.083 | 0.011 |
| Mean |  |  |  |  |  |
| B | 6.4 | 0.64 | 0.64 | - | 0.018 |
| O | 9.8 | 0.60 | 0.62 | - | 0.033 |

P-value calculated by using a Markov chain, performed in Arlequin 3.5.2.2 B- Breeders (n=23); O- Offsprings (n= 244); N<sub>a</sub>, Allele number; H<sub>o</sub>, Observed Heterozygosity ; H<sub>e</sub>, Expected Heterozygosity; P<sub>H-W</sub>, P-values for Hardy-Weinberg equilibrium; F<sub>IS</sub>, Inbreeding coefficient . \*P<0.01; \*\*P<0.001.

After analyzing the mother’s genotype in each spawn, it was observed that some offspring did not corresponded to the mother. This happened because octopus hatchlings are able to *escape* from their original incubator and *jump* into another one. These hatchlings, together with the samples with undetectable signals in the fragment analysis, were excluded from the parentage analysis. Fathers were assigned to 244 octopus juveniles for which the mother was known. The results obtained with GERUD and COLONY revealed evidence of high levels of multiple paternity in all analyzed spawns.

The estimated minimum number of sires from the GERUD analyses ranged from three to five, with an average of 4.4 sires per spawn (Table 6, Fig. 3A). The mean maximum number of sires estimated with COLONY was 10.2 per spawn.

**Table 6.**
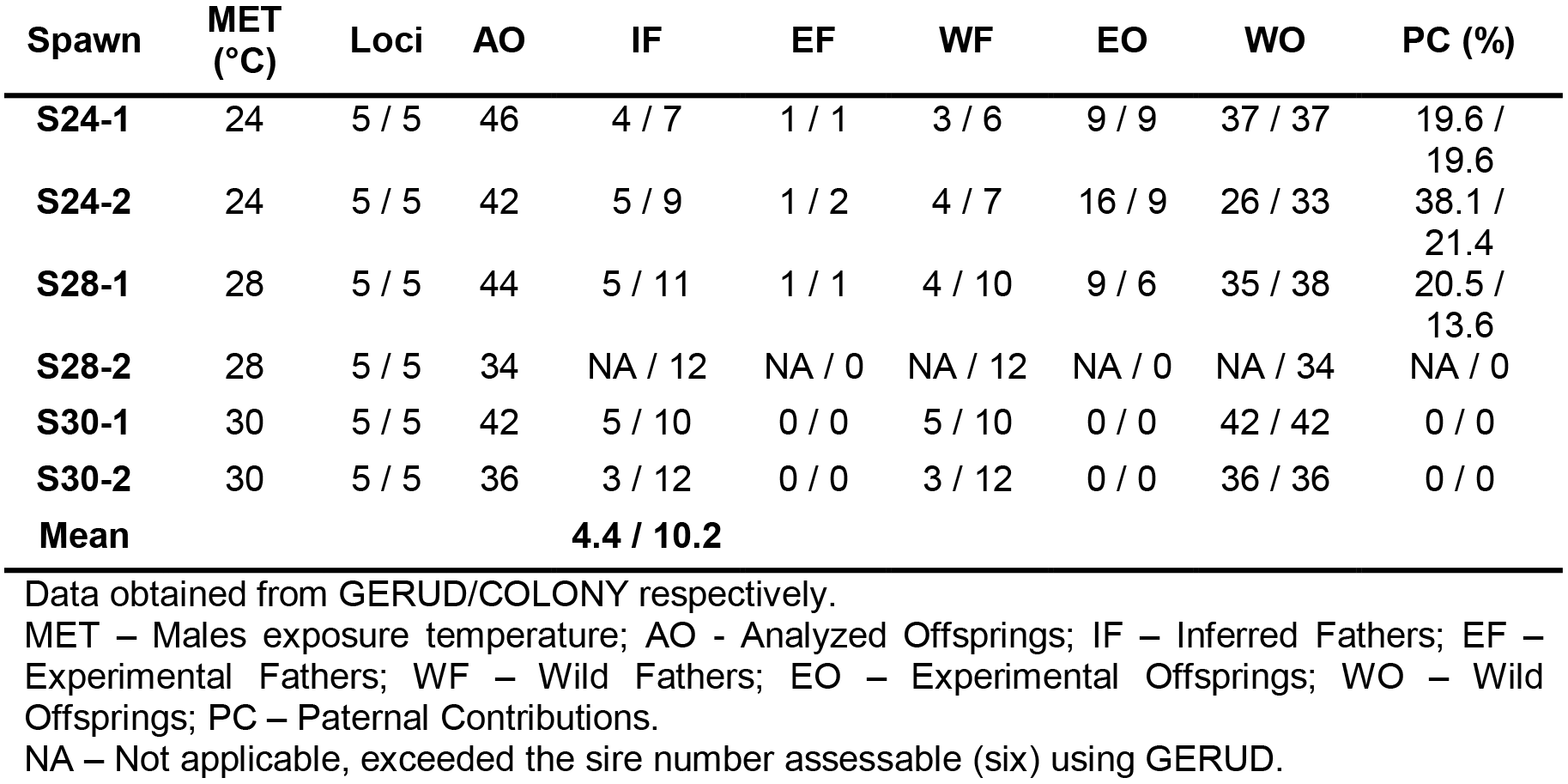
Number of sires assigned with the paternal analysis for each spawn of *O. maya* using COLONY and GERUD.

**Fig. 3.**
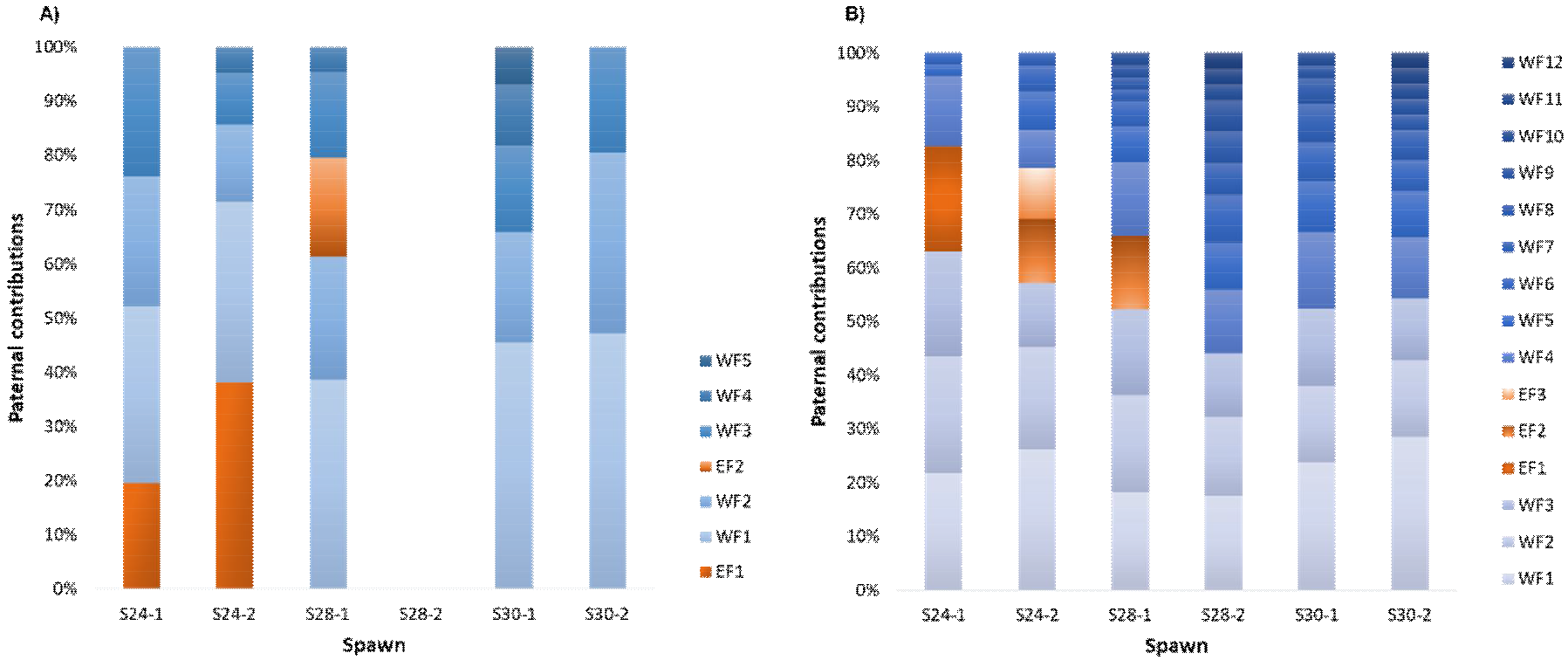
Relative contributions of sires in each spawn of *Octopus maya* using GERUD (A) and COLONY (B). EF - Experimental fathers (1-first male mated; 2-Second mated and 3-Third); WF1-WF12: All unknown wild fathers.

According to the parentage analysis using GERUD, when the males were exposed at 24°C, only one experimental male contributed to the progeny (S24-1 and S24-2 in both the 1^st^ male; Table 6, Fig. 3A); these males were the sires of nine and 16 offsprings, with a contribution of 19.6% and 38.1%, respectively, of the analyzed progeny. When the males were exposed at 28°C, one experimental male was identified as potential sire of 9 offspring (S28-1 the 2^nd^ male), contributing with 20.5% of the analyzed progeny (Table 6, Fig. 3A). In the case of the spawn S28-2 paternity could not be assigned. When males were acclimated to 30°C, they had no contribution to the progeny (S30-1 and S30-2), but the minimum number of sires were 5 and 3, respectively. It was assumed that under this experimental condition all progeny belongs to wild males (Table 6, Fig. 3A).

The COLONY analysis results showed that when the males were exposed at 24°C for 30 d, one to two experimental males contributed to the progeny with 19.6% and 21.4%, respectively, of the analyzed progeny (S24-1 the 1^st^ one with 9 offspring, and S24-2 the 2^nd^ and 3^rd^ male with 5 and 4 offspring, respectively; Table 6, Fig. 3B). When the males were exposed at 28°C, one experimental male was identified with 6 offspring and a parental contribution of 13.6% (S28-1). In the S28-2 spawn no sires were identified (Table 6, Fig. 3B). Males exposed at 30°C, showed no contribution to the progeny. It was assumed that all the offspring correspond to wild male octopuses.

The COLONY analysis results also showed that independently of the maximum number of sires that explains the progeny, there are at least four males which contributed with the 57.1% of the total progeny per spawn, and the other 42.9 % is distributed among the remaining parents (Fig. 3B).

## Discussion

Previous studies showed that temperature modulates the reproductive capacity of *O. maya* wild populations, reducing the functional maturity and SCI (%) when environmental temperature in the YP continental shelf is around 30°C (Angeles-Gonzalez et al., 2017). The present study was designed to evaluate if temperatures higher than 27°C affect the reproductive capacity and success of male *O. maya* as observed when females and their embryos were exposed to thermal stress (Juárez et al., 2015; Juárez et al., 2016; Sanchez-García et al., 2017). Results obtained in the present study, demonstrate that temperature of 30°C affected negatively growth rate. For the digestive gland index of the adult *O. maya* males a negative effect was observed at temperatures from 28°C to 30°C. *O. maya* males exposed to 30°C showed intermittent feeding, possibly as a consequence of the exposure to high temperatures, as reported in *O. pallidus* (André et al., 2008). The deleterious effect of the temperature on the digestive gland could directly affect the reproductive performance because most of the energy that is directed to reproduction comes from this organ. At the same time, an increment of haemocytes, and a reduction on VO_2_ were registered, indicating that several physiological mechanisms were affected in this thermal condition. In mollusks, in the absence of a specific immune system the immune response is mediated by circulating haemocytes and molecular effectors that allow a rapid and effective response to stressors. In bivalve mollusks such as *Chamelea gallina* exposed to 30°C, and cephalopods such as *Eledone cirrhosa* it was observed an increment in the circulating haemocytes (THC) when the organisms were exposed to different stressors, as observed in *O. maya* males (Malham et al., 2002; Monari et al., 2007).

Octopuses are aquatic ectotherms, an increment in temperature provokes an increment in the energetic demands that are essentially covered in first instance to maintain the homeostasis, even if the cost reduces growth (Sokolova et al., 2012). In adult *O. maya* males a reduction of the oxygen consumption and growth jointly with a decrease on DGI (%) was observed in animals maintained at 30°C. In *Sepia officinalis* it was observed that the oxygen consumption of animals from the English channel acclimated to 21°C showed a metabolic rate lower than observed in cuttlefish acclimated to 15°C (Oellermann et al., 2012). That pattern of thermal acclimation was explained by taking into account that a suppression of oxygen consumption rates in organs other than the hearts (e.g. digestive gland, mantle, or even reproductive tissues) could be occurring in this species. Although the tissue oxygen consumption was not measured in this study, we can hypothesize that as in cuttlefish, in *O. maya* there are compensatory mechanisms that reduce food ingestion and digestive gland metabolism to save energy, allowing the key organs such as the heart, to maintain the homeostasis of the animal, at least temporarily.

From a reproductive point of view, the 30°C temperature treatment affected various levels of the testis organization: dilation of seminiferous tubes, shrinkage of the proliferative stratum where spermatozoa are synthetized, high quantity of acidophilic bodies, and a general disorder in the organization of the germinal tissue. Although temperatures higher than 27°C affected the reproductive efficiency of this species (Juárez et al., 2015; Juárez et al., 2016; Sanchez-García et al., 2017), this is the first time that the effects of temperature on the reproductive capacity at histological level of adult males are reported, demonstrating that a temperature of 30°C strongly restricts the reproduction of males in this species (Angeles-Gonzalez et al., 2017).

Temperature of 30°C affected the structures of reproductive tissues in the adult males, provoking an inflammatory process in the testis and a higher disorder at the tissues than that observed in animals maintained at 24°C. An intermediate condition was observed in animals maintained at 28°C, suggesting that this may be a thermal threshold for reproduction of male *O. maya*. While temperature did not affect the number of spermatozoa per spermatophore, a higher production of spermatophores was observed in animals maintained at 30°C. This suggests that despite the structural damage caused by temperature, animals responded by allocating enough energy to increase their reproductive potential. This could be a reproductive strategy to ensure the preservation of the species, through the formation of a greater number of spermatophores. Although we don’t know if there is a direct relationship between quantity of live sperms and fertilization rate in *O. maya,* it is possible to think that a higher GSI could be activated as a compensatory mechanism to reduce the effects of changes in the testis structure due to thermal stress, increasing the fecundity probability of thermal stressed animals (Parker, 2016).

The analysis of six spawns with five different microsatellite loci in the progeny of six females confirmed the presence of multiple paternity in *O. maya*. A minimum number of four and a maximum of 10 males were estimated to contribute to the progeny. This conserved reproductive strategy has been observed in other octopod species such as *Graneledone boreopacifica* (Voight and Feldheim, 2009), *Enteroctopus dofleini* (Larson et al., 2015), *O. vulgaris* (Quinteiro et al., 2011) *O. oliveri* (Ylitalo-ward, 2014) and *Euprymna tasmanica* (Squires et al., 2014). It was also observed that the last mated experimental male had no parental contribution in any spawn, with exception of male S24-2, whose parental contribution was lower than that of the other males involved. Contrary to the pattern of the last male precedence observed in *Loligo bleekeri* (Iwata et al., 2005), in *O. maya,* the last male to copulate is not the best genetically represented in the offspring. The pattern identified in *O. maya* coincides with the pattern of first male precedence observed in *O. oliveri* (Ylitalo-ward, 2014). Indeed, under optimal conditions (24°C) the experimental males contributed with an average of 57.1% of the total parental contribution for each spawning, regardless of the order of mating. However, several studies have shown that spermatic precedence is influenced by the order of mating, due to sperm competition, or mediated by female cryptic choice (Iwata et al., 2005; Quinteiro et al., 2011; Hirohashi and Iwata, 2016). This apparent disagreement may reflect the high diversity in cephalopod reproductive strategies.

Temperature increases plays an important role in the parental contribution (reproductive success) of *O. maya* due to the fact that in the spawning of stressed parents (28°C) a reduction in the parental contribution was observed. This was more evident at 30°C where no contribution of the experimental males was found, independent of the mating order.

Temperature affected the growth and the metabolism of *O. maya* males by reducing the food ingested and the digestive gland index; as a consequence, the organism directed available energy to reproduction. Males under stress conditions produced a greater number of spermatophores. Nevertheless, this strategy seems to be insufficient given the testis damage at high temperatures. Both, paternity and histological analyses showed that the 28-30°C thermal range affects the reproductive success of *O. maya* adult males, independently of the compensatory mechanisms activated in response to the damage.

Results obtained in the present have demonstrated that temperature is a strong environmental factor that determines the reproductive success of *O. maya,* both in laboratory and in wild populations (Juárez et al., 2015; Angeles-González et al., 2017). In some cephalopod species studies, data demonstrate that temperature higher than experienced in wild conditions, can shorten the period of sexual maturity, reducing it by half (Takahara et al., 2016). Although this response could be apparently advantageous allowing the proliferation of cephalopods around the world (Doubleday et al., 2016), results obtained in this study evidence that in this species males and females have a temperature threshold for reproduction around 28°C, above of which the physiological condition, the reproductive performance and success are significantly reduced.

## Acknowledgments

The present study was done at the Laboratory of cephalopod production in UMDI-UNAM, Sisal Yucatan under financial support of DGAPA-PAPIIT Program IN219116 from Universidad Nacional Autónoma de México. All genetic analyses were done at Functional Marine Genomic Laboratory, at the department of Marine Biotechnology in Centro de Investigación Científica y Educación Superior de Ensenada, México (CICESE).

Research was supported by the SEP-CONACYT-CB-2014-01/241690 grant.

We would like to thank CONACYT and CICESE for the scholarship granted to Laura López-Galindo; the results presented here are part of her Doctoral Dissertation at CICESE.

We thank M.C. José F. Tercero for octopus capture and laboratory conditioning; Zoila Peregrina Canté Cuá, Ricardo Salomone Lopes, Karina Nambo-García and Itzel Tapia for sampling. This paper is part of the ‘TempOxMar’ collaboration research net (Evaluación de los efectos de la temperatura y el oxígeno disuelto en poblaciones de organismos bentónicos marinos de interés pesquero, ecológico y acuícola) organized by Universidad Nacional Autónoma de México (UNAM) and supported by Dirección General de Internacionalización-UNAM. A. Olivares is grateful for the sabbatical year (2015-2016) provided by Universidad de Antofagasta, Chile, during which this work was developed. Avila-Poveda OH is commissioned as CONACYT Research Fellow/UAS-FACIMAR (Project No. 2137), and participated as a member of the academic group ‘Manejo de Recursos Pesqueros UAS-CA-132, UAS-FACIMAR’ accredited to ‘TempOxMar’ and obtained research residency at UNAM under the Annual Program of Academic Cooperation UAS-UNAM (2016-NI-0036A001P001/02/03).

## Competing interests

No competing interests declared.

## Funding

This research was founded by projects PAPIIT IN219116 to CR, SEP-CONACYT-CB-2014-404 01/241690 and CICESE: 682123 to CEG.

## Author contributions

L.L.G., C.G.S, C.R. designed the experiments; L.L.G., C.G.S., C.R. A.O., O.H.A.P., wrote and revised the paper; L.L.G., and Z.C.C. conducted animal experimental management and care procedures; L.L.G., Z.C.C., C.R., F.D. conducted physiological assessments; L.L.G., Z.C.C, C.R., F.D. performed dissection, sampling and sperm quality assessments; L.L.G, C.R., A.O., O.H.A.P. performed the histological analysis; L.L.G. performed the statistical analysis; L.L.G., J.P.P., K.N.G., O.E.J. DNA extractions; J.P.P., K.N.G., O.E.J., F.L.C. performed microsatellite amplification; L.L.G., J.P.P., K.N.G., O.E.J., F.L.C., C.G.S. Genotype assignment; L.L.G., J.P.P., O.E.J, C.G.S. Parentage analysis; C.G.S., C.R., F.L.C supplied materials and supervised methodology.

---

**Table S1.**
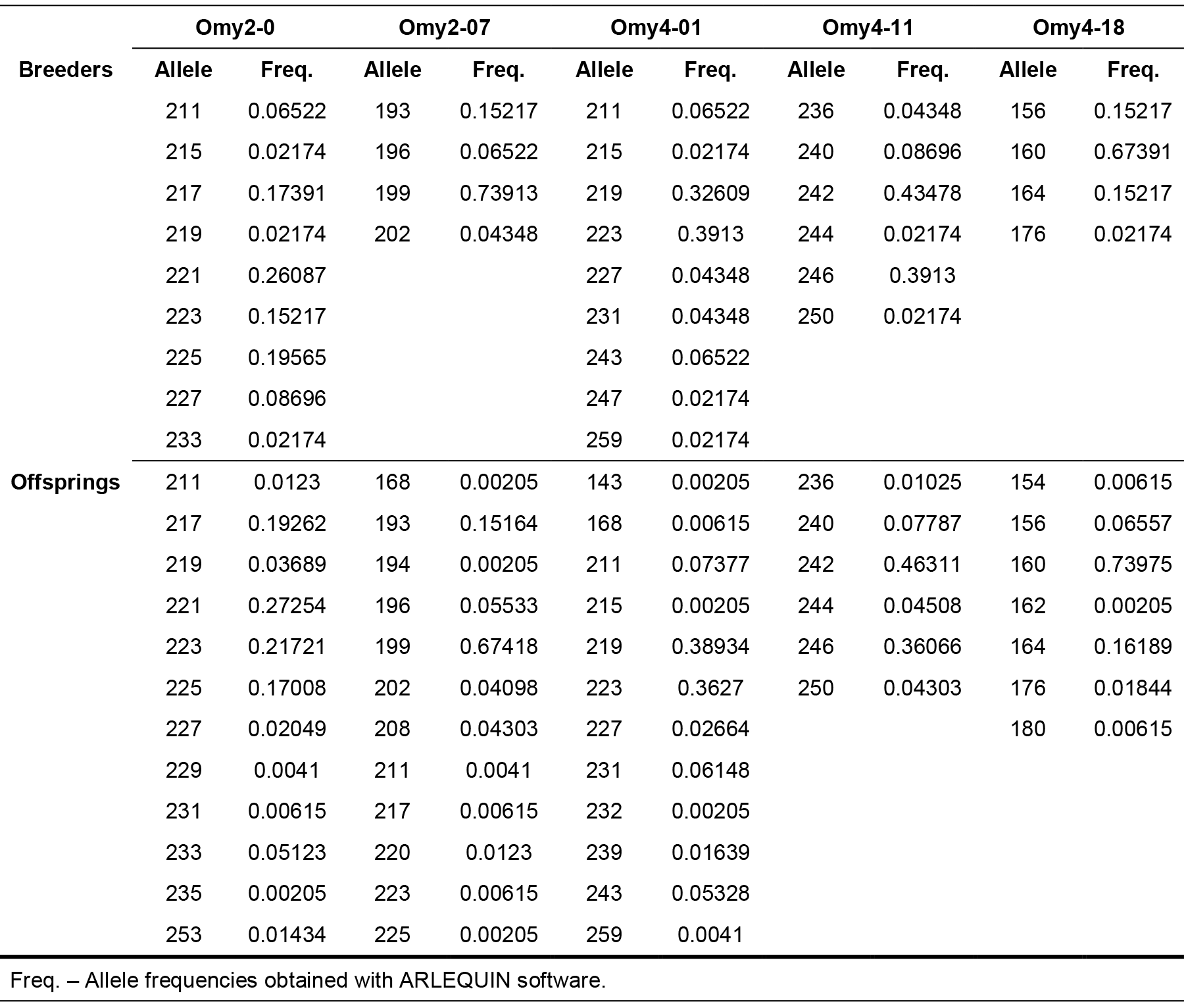
Allele frequencies for each microsatellite locus of *O. maya* males.

| Breeder | Omy2-0 |  | Omy2-07 |  | Omy4-01 |  | Omy4-11 |  | Omy4-18 |  |
| --- | --- | --- | --- | --- | --- | --- | --- | --- | --- | --- |
|  | Allele | Freq. | Allele | Freq. | Allele | Freq. | Allele | Freq. | Allele | Freq. |
| Breeder | 211 | 0.06522 | 193 | 0.15217 | 211 | 0.06522 | 236 | 0.04348 | 156 | 0.15217 |
|  | 215 | 0.02174 | 196 | 0.06522 | 215 | 0.02174 | 240 | 0.08696 | 160 | 0.67391 |
|  | 217 | 0.17391 | 199 | 0.73913 | 219 | 0.32609 | 242 | 0.43478 | 164 | 0.15217 |
|  | 219 | 0.02174 | 202 | 0.04348 | 223 | 0.3913 | 244 | 0.02174 | 176 | 0.02174 |
|  | 221 | 0.26087 |  |  | 227 | 0.04348 | 246 | 0.3913 |  |  |
|  | 223 | 0.15217 |  |  | 231 | 0.04348 | 250 | 0.02174 |  |  |
|  | 225 | 0.19565 |  |  | 243 | 0.06522 |  |  |  |  |
|  | 227 | 0.08696 |  |  | 247 | 0.02174 |  |  |  |  |
|  | 233 | 0.02174 |  |  | 259 | 0.02174 |  |  |  |  |
| Offspring | 211 | 0.0123 | 168 | 0.00205 | 143 | 0.00205 | 236 | 0.01025 | 154 | 0.00615 |
|  | 217 | 0.19262 | 193 | 0.15164 | 168 | 0.00615 | 240 | 0.07787 | 156 | 0.06557 |
|  | 219 | 0.03689 | 194 | 0.00205 | 211 | 0.07377 | 242 | 0.46311 | 160 | 0.73975 |
|  | 221 | 0.27254 | 196 | 0.05533 | 215 | 0.00205 | 244 | 0.04508 | 162 | 0.00205 |
|  | 223 | 0.21721 | 199 | 0.67418 | 219 | 0.38934 | 246 | 0.36066 | 164 | 0.16189 |
|  | 225 | 0.17008 | 202 | 0.04098 | 223 | 0.3627 | 250 | 0.04303 | 176 | 0.01844 |
|  | 227 | 0.02049 | 208 | 0.04303 | 227 | 0.02664 |  |  | 180 | 0.00615 |
|  | 229 | 0.0041 | 211 | 0.0041 | 231 | 0.06148 |  |  |  |  |
|  | 231 | 0.00615 | 217 | 0.00615 | 232 | 0.00205 |  |  |  |  |
|  | 233 | 0.05123 | 220 | 0.0123 | 239 | 0.01639 |  |  |  |  |
|  | 235 | 0.00205 | 223 | 0.00615 | 243 | 0.05328 |  |  |  |  |
|  | 253 | 0.01434 | 225 | 0.00205 | 259 | 0.0041 |  |  |  |  |

Freq. – Allele frequencies obtained with ARLEQUIN software.

**Fig. S1.**
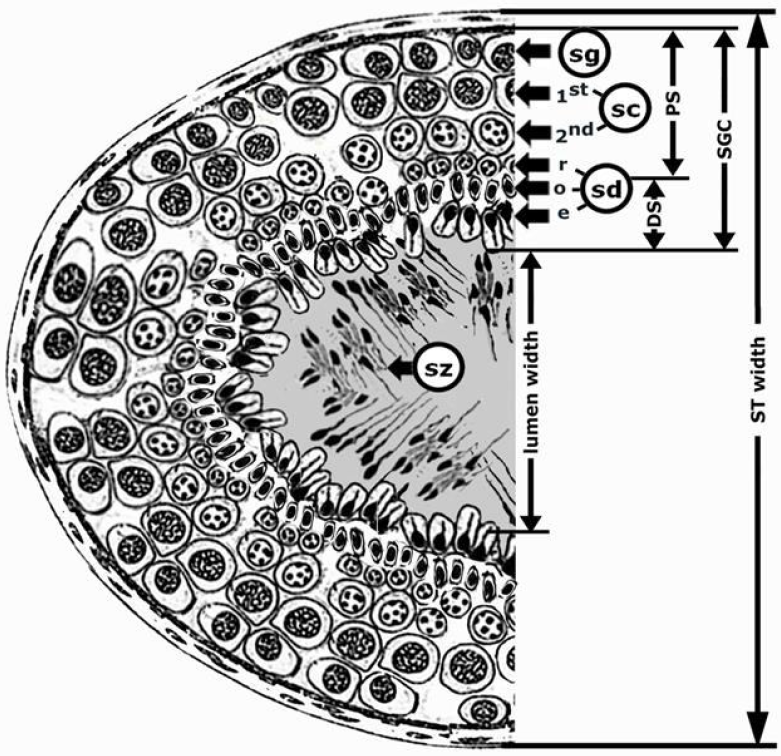
Schematic drawing of one seminiferous tubule (ST) in longitudinal section showing measured widths and heights. Abbreviations: DS-differentiation stratum, PS-proliferative stratum, SGC-strata of germ cells(SGC = PS + DS), sg-spermatogonia, sc-spermatocytes, sd-spermatids (round, ovoid and elongated), sz-spermatozoa. The grey shadow represents the lumen where sperm is free (spermiation).

**Fig. S2.**
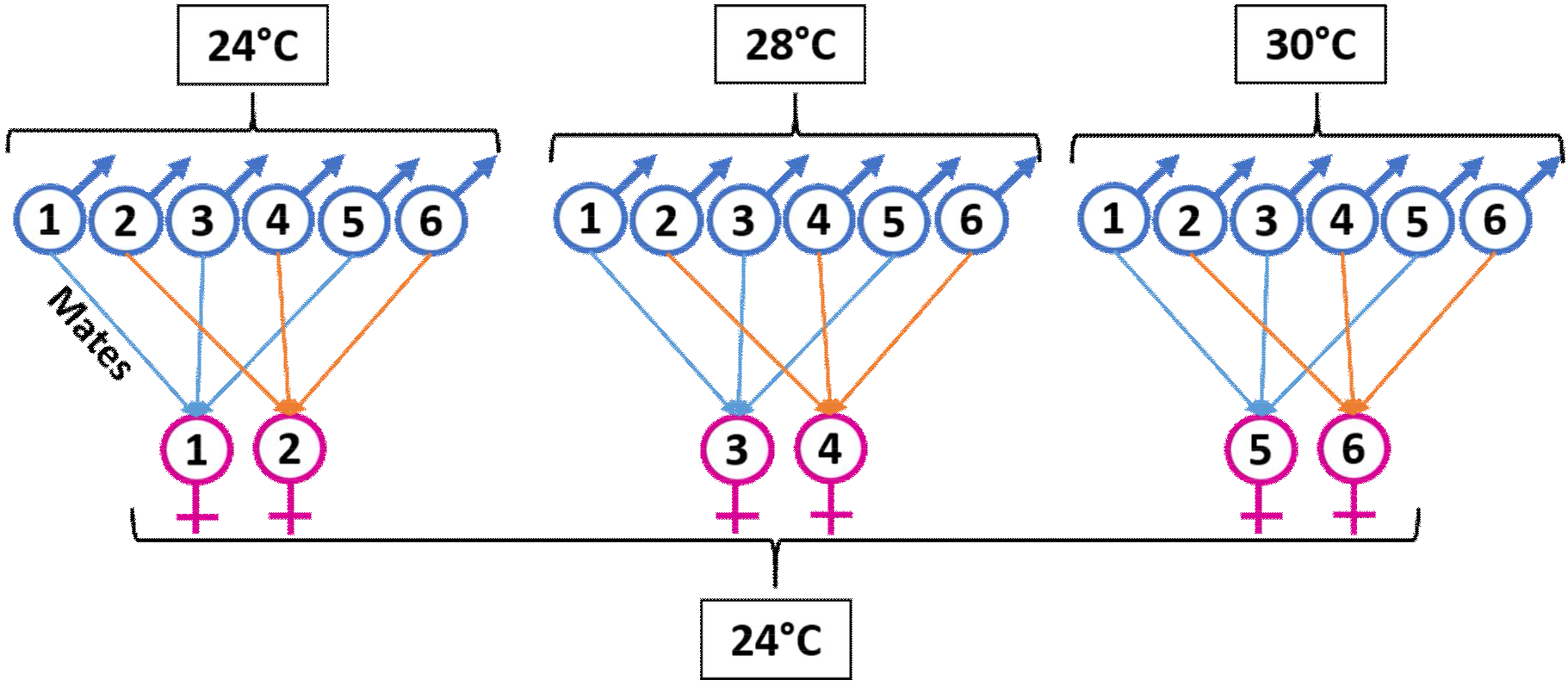
Mating system used in *Octopus maya* per experimental temperature. Males maintained at different experimental temperatures (24°C, 28°C and 30°C) during 30 d were mated with females at 24°C. The matings were done one by one for each temperature. Copulation lasted 4 to 6 hours and between each mating the females had a recovery time of 4 d until the next mating. The females were acclimated for 15 d at 24°C until mating.

## List of abbreviations

°C: Degrees Celcius
μl: microliters
μM: Micromolar
μm: Micrometres
Abs: Absorbance
ASP: Percentage of alive spermatozoa
B: Breeders
BW: Octopus total body weight
cm: Centimetres
d: Days
DF: Dilution factor
DGI: Digestive gland index
DGW: Digestive gland weight
DO: Dissolved oxygen
DS: Differentiation stratum
ε: Extinction coefficient
F: Water flow rate
F_IS_: Inbreeding coefficient
g: Grams
GSI: Gonadosomatic index
h: Hours
Hc: Hemocyanin concentration
H_o_/H_e_: Observed/Expected heterozygosity
hOP: Hemolymph osmotic pressure
H_W-E_: Hardy-Weinberg equilibrium
L: Litres
L/D: Light/Dark
Ln: Natural logarithm
m: Meters
mg: Milligrams
min: Minutes
ml: Millilitre
mM: Millimolar
mm: Millimetres
n: Specimens number
Na: Allele number
ng: Nanograms
O: Offsprings
*O. maya*: Octopus maya
O_2i_ / O_2o_: Oxygen concentration of the water inlet/outlet
OP: Osmotic pressure
OsmC: Osmoregulatory capacity
P: Probability
PCR: Polymerase chain reaction
pH: Hydrogen potential
ppt: Parts per thousand
PS: Proliferative stratum
PVC: Polyvinyl carbonate
SCI: Spermatophoric complex index
SCW: Spermatophoric complex weight
SD: Standard deviation
SGC: Strata of germ cells
SGR: Specific growth rate
ST: Seminiferous tubules
STN: Spermatophores total number
t: Time
TASC: Total number of alive spermatozoa
THC: Total haemocytes count
TSC: Total Number of spermatozoa
TW: Testis weight
VO_2_: Oxygen consumption
WG: Weight gain
Wi/Wf: Initial/Final weight
wOP: Water osmotic pressure
ww: Wet weight
YP: Yucatan Peninsula

